# CAFT: A Compositional Log-Linear Model for Microbiome Data with Zero Cells

**DOI:** 10.1101/2025.11.26.690468

**Authors:** Glen A Satten, Mo Li, Ni Zhao

## Abstract

**Background:** Differential abundance analysis is fundamental to microbiome research and provides valuable insights into host-microbe interactions. However, microbiome data are compositional, highly sparse (with many zero counts), and influenced by differential experimental biases across taxa. Standard statistical methods often overlook these features. Many approaches analyze relative abundances without accounting for compositionality or rely on pseudocounts, potentially leading to spurious associations and inadequate false discovery rate (FDR) control.

**Methods:** We introduce a novel framework for differential abundance analysis of microbiome data: the Compositional Accelerated Failure Time (CAFT) model. CAFT addresses zero read counts by treating them as censored observations that are below a detection limit. This approach is inherently resistant to multiplicative technical bias, eliminates the need for pseudocounts, and addresses compositional bias through the establishment of appropriate score test procedures.

**Results:** Extensive simulations show that CAFT outperforms competing compositional differential abundance methods, including LOCOM, LinDA, ANCOM-BC2, its robust variant, and LDM-clr by offering more robust type I error and FDR control with or without technical bias. Additionally, we applied CAFT to microbiome data on inflammatory bowel disease (IBD) and the upper respiratory tract (URT) to identify differentially abundant gut microbial taxa between IBD patients and healthy controls, as well as URT taxa distinguishing smokers from non-smokers.

**Conclusion:** We present CAFT, a powerful, robust, and efficient approach for compositional differential abundance analysis. CAFT effectively controls Type I error and maintains FDR control, while demonstrating enhanced power in statistical testing. These capabilities render CAFT a useful tool for compositional microbiome data analysis.

**Availability and implementation:** The R package and Vignette are available at https://github.com/mli171/CAFT.

## Introduction

High-throughput sequencing technology has revolutionized our understanding of biological processes and their impact on human health. In many instances, initial experimental outcomes from these techniques were obtained before the establishment of robust statistical methods for their analysis. Subsequently, statistical methods tailored to the characteristics of each data type are developed, which bring the ability to produce replicable results. This, coupled with standardized experimental procedures, ensures that the observed signals reflect biological effects rather than experimental artifacts. Microbiome sequencing studies may eventually follow this path as well. Currently, microbiome studies suffer from a lack of methods to standardize experimental results. This is largely because the abundance measurement of each taxon may be affected by different amounts of bias (1). However, even if ‘model community’ samples with known relative abundances are included as experimental controls, we may only be able to learn about the bias of the (small number of) taxa in the model community. While it may be possible someday to use model community data to infer a standardization for *every* taxon in a sample (2), this approach is not currently viable, and in particular, data on the bacterial and genomic factors that affect bias have not been studied.

Fortunately, there are steps that can be taken at the data analysis stage to account for the lack of taxon-specific standardization in microbiome data. A recent model (1) suggests that the bias in taxonomic counts can be represented by multiplicative factors, a result later validated by (2). An important consequence of this is that unbiased estimates of key parameters in carefully constructed models can still be obtained even from data that are subject to the experimental bias described by this model. For example, unbiased estimates of odds ratios comparing two taxa can be obtained using logistic regression even if data have not been standardized to account for biases, as implemented in LOCOM (3).

Although logistic regression can estimate odds ratios, most compositional microbiome data analyses use log-linear models. These models typically utilize log-transformed count or relative abundances, effectively addressing the compositional constraint of uninterpretable total counts across all taxa. Further, differences in regression coefficients across taxa or, equivalently, constraining regression coefficients to sum to zero ((4), (5)), estimate parameters that are like log-odds ratios, and hence are bias-robust. However, this method faces major challenges in microbiome analyses due to the prevalence of zero counts. Most compositional methods deal with zeros by adding a *pseudocount* (e.g., 1, 1/2, or another small value) either to the zero cells, or to all cells (6–9). Unfortunately, the choice of pseudocount can affect results, especially if the number of zero cells relates to the variable of interest. Further, the pseudocount does not scale with the overall taxon abundance, leading to biased estimates even when the experiment is free of technical bias. The only current approach that rigorously handles zero counts is LOCOM (3), which avoids log-transformations by using logistic regression.

Here, we introduce a novel approach to analyzing compositional microbiome data by treating zero counts as values that fall below a sample-specific limit of detection or, equivalently, as *censored observations* in survival analysis. Similar strategies have been successfully applied in other settings with data below a detection limit (10–13). In this work, we consider the negative log-relative abundance as a right-censored survival time. The censoring time is the log of the library size. Thus, fitting a log-linear model for microbiome data becomes equivalent to fitting an accelerated failure time (AFT) model. This is consistent with the pseudocount approach that the zero cell is treated as a ‘small’ value, but ensures that the data for each taxon scales appropriately. We note that (14) also treats zero cells as censored observations, but uses a Cox proportional hazards model to relate covariates to relative abundance. This model, however, is not compositional and thus is not invariant to bias. Its main contribution is to replace the two-sample Wilcoxon test sometimes used to test the effect of a binary covariate on relative abundances of a single taxon with the log-rank test (which, as the score test for the Cox model, allows for more general covariates). Since the approach is not compositional, a log transformation is not necessary, and the hypothesis tested by this model is in fact that tested by the Linear Decomposition Model (15).

The AFT is often fit using parametric models; this is only appealing if a reasonable choice of distribution can be made. Alternatively, there are rank-based methods that do not require a distributional choice. Here we use the rank-based approach of (16) and (17), which we have found to have good behavior even when the proportion of censored data is very high. We reformulate the rank-based score equations into a convex Jaeckel-like dispersion (18) or objective function, which is then minimized to obtain the model parameter estimates. Inference is based on score tests, since the rank-based score is a step function, so its ‘derivative’ (required for Wald-like tests) requires smoothing. The ‘usual’ estimator of the variance of the score function is based on a connection with the Cox proportional hazards model; instead, we use a novel, simpler estimator of the score function variance based on rank methods that we find outperforms the standard estimator. Finally, by using rank-based inference, we hope to develop asymptotic tests of the hypotheses of interest, rather than the Monte-Carlo methods needed for LOCOM. For this reason, we also avoid the Monte-Carlo variance-covariance estimator of the estimated parameters of (19).

## Methods

The fundamental idea behind CAFT is to use the AFT to fit a log-linear model to relative abundance from each taxon, then base inference on differences between regression parameters. This approach avoids pseudocounts, but still encounters the problem of how to estimate the sample-specific mean of the log relative abundance (which corresponds to the normalization constant) when some taxa have zero counts. We next show how treating this value as a random effect allows a simple solution.

### The log-linear model for microbiome data

Our starting point is the log-linear model for microbiome data in the presence of experimental bias proposed by (1) and extended by (2) to include covariates. We let *C*_*ij*_ denote the counts of taxon *j* in sample *i, N*_*i*_ = ∑_*j*_ *C*_*ij*_ denote the library size. We adopt the notation that for matrix *M, M*_*i·*_ corresponds to the row vector that is the *i*th row of *M* while *M*_*·j*_ is the column vector that is the *j*th column of *M*.

Let 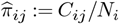 represent the observed relative abundance of taxon *j* in sample *i* and let 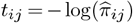 with the understanding that *t*_*ij*_ = ∞ when 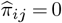 zero cell corresponds to *C*_*ij*_ *<* 1, i.e. 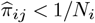. Thus, we may consider 1*/N*_*i*_ to be the ‘limit of detection’ for observing relative abundances. Equivalently, we may consider the values of *t*_*ij*_ to be censored whenever *t*_*ij*_ *>* log(*N*_*i*_). Thus, we define a ‘censored’ version of *t*_*ij*_, denoted *τ*_*ij*_ and defined by 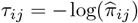 if 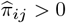 and if 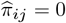. The corresponding censoring indicator is 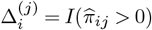. For each taxon *j*, then, a log-linear model for 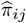 values corresponds to the AFT model

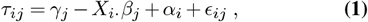

which is the censored-data version of a log-linear model. In equation Eq. (1), *X* is a matrix with *i*th row comprising the covariates of the *i*th observation, *β*_*j*_ is a vector of regression parameters that describe the effect of covariates *X* on the log-relative abundances, *γ*_*j*_ is a taxon-specific intercept and *E*_*ij*_ is residual error. Because an intercept is explicitly included, *X* should not contain an intercept (constant) column, and the columns of *X* must be centered (i.e., sum to zero). The term *α*_*i*_ plays the role of a normalization constant, and is here treated as a random effect because centering *τ*_*ij*_ by its mean over taxa is problematic with censored data. The taxon-specific intercept *γ*_*j*_ can be written as *γ*_*j*_ = −*ln*(*π*_*j*_) + *ω*_*j*_ where *ln*(*π*_*j*_) is the ‘true’ value of the log-relative abundance (i.e., the expected value of log-relative abundance we would obtain in an unbiased experiment) for a sample having *X*_*i·*_ = 0, and *ω*_*j*_ is the ‘bias factor’ of taxon *j* as introduced by (1). Here we treat *γ*_*j*_ as a nuisance parameter.

The presence of the random effect *α*_*i*_ is problematic for rank-based analysis. To see how we can deal with this, we first suppose that there were no zero cells, i.e., that there were no censored observations. If the *α*_*i*_ values were known, we could fit Eq. (1) using ordinary least squares. However, *α*_*i*_s are unknown, and if we naively ignore *α*_*i*_s, we will obtain a biased estimate of the *β*_*j*_s if the random effect has any correlation with covariates *X*. In particular, if there were no censored observations, we would find that by ignoring the random effect, we had estimated 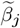 given by

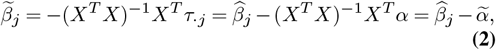

where 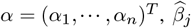 is the ‘correct’ estimate, and 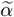 is the error we make in estimating *β*_*j*_ while ignoring the random effect. Note however that this bias term is the same for each taxon *j*. Thus, we can estimate *β*_*j*_ − *β*_*j’*_ by the estimator 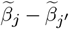 since this estimator is identically equal to 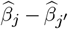 for least-squares regression with no censored data. Fortunately, 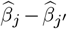 corresponds to an odds ratio, which has been shown to be a parameter that can be identified free of bias (1, 3).

In the presence of censoring (i.e., zero cells), we cannot use ordinary least squares to fit Eq. (1). Although the Buckley-James (20) estimator attempts a least-squares-like approach, its estimating equation is known to have multiple roots. Thus, we propose to use the rank-based estimator of *β*_*j*_ from (16) and (17), which is known to have a single solution and is robust to outlying values of the dependent variable. Because these estimators converge asymptotically to the same parameters as in the linear model with complete data, we can ignore the part of the random effect *α*_*i*_ that is along *X* when we fit regression parameters *β*_*j*_, so long as we confine our interest to differences *β*_*j*_ − *β*_*j’*_. (The part of the random effect *α* that is orthogonal to *X* can be considered part of the random error *ϵ*.)

We estimate *β*_*j*_ by solving the following score equations:

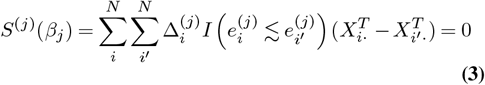

where 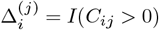 is the censoring indicator, 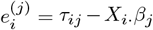 and 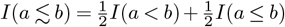. We note that equation Eq. (3) can be rewritten in a more compact form similar to the score function for rank-based regression (i.e., *R*-estimation) in (21) as

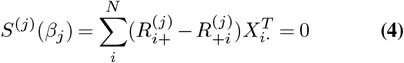

where 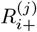 and 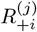 are the row and column sums, respectively, of the matrix 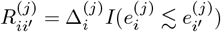. Given representation Eq. (4), it is also possible to show that *β*_*j*_ can be estimated by minimizing a Jaeckel-like dispersion

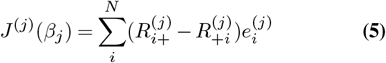

which, like the Jaeckel dispersion (21), can easily be shown to be convex and is in fact equivalent to the objective function in (17).

Equations Eq. (3) and Eq. (4) do not depend on the intercept *γ*_*j*_. If an estimator of *γ*_*j*_ is desired, one could calculate the Kaplan-Meier estimator of the values 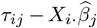 using the censoring indicator Δ_*ij*_ and then estimate *γ*_*j*_ by the median obtained from the Kaplan-Meier estimator.

### Testing hypotheses about regression coefficients

As only differences in *β*_*j*_s are interpretable, it is necessary to make additional assumptions to test biologically-meaningful hypotheses. If we could identify taxa that we were sure were ‘null’ (i.e., had relative abundance that did not vary with covariates *X*), we could make our comparisons to these taxa. As this information is generally unknown, we adopt the assumption that far fewer than half of the taxa are non-null, following the methodology described in (3) and (7). Consequently, we use the median value of 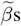, denoted by 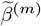, as the null value in our hypothesis tests. (We note that the modal value, used by (6) and available in (7), is also an attractive choice). To test whether covariates *X* affect the relative abundances of taxon *j* under the assumptions described in the previous section, we must test hypotheses like 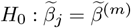 where 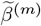. In general, this median (or mode) is unknown and should contribute to the variability of the test. However, since 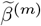 is estimated using data from *all* taxa, we take 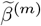 to be a fixed parameter. If we are testing a hypothesis with *d >* 1 components, we take 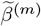 to be the affine equivariant spatial median (22) as implemented in the R function `HR.Mest` in the `ICSNP` package.

The variance-covariance matrix of 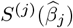 based on the Martingale representation of the Cox model is given by (19). We denote this variance estimator as the Cox estimator of the variance of the score function. Because of some typographic errors in (19), we give the expression for the Cox variance estimator in Section S1 of the Supplementary Materials. Alternatively, representation Eq. (4) also suggests a simple variance estimator for the score function 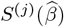, as it implies independence between the rank information in *R* and the residuals *e* at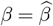. If we let *V*^(*j*)^ denote the variance-covariance matrix of *S*^(*j*)^, the estimator 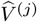 of *V*^(*j*)^ implied by Eq. (4) is

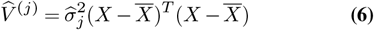

where

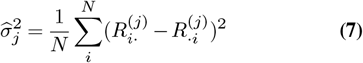

and 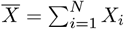. We refer to this variance as the Rank estimator of the variance of the score function.

The main drawback of score tests is that we require a restricted estimator of nuisance parameters not specified under the null. If *β* has *K* components and the (composite) null hypotheses specify only *d < K* of these, then assume that we wish to test hypotheses of the form

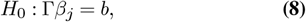

where Γ is a *d × K*-dimensional matrix and where we assume Γ has full row rank. We let Λ be any matrix having orthogonal rows that span the null space of Γ, i.e. Λ is a (*K* − *d*) *× K*-dimensional matrix that satisfies ΓΛ*T* = 0_*d*,(*K*_ − _*d*)_. For example, if *K* = 3 and we wish to test if the first two components are equal, we could choose Γ = (1, −1, 0), *b* = 0 and then take 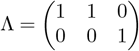. Note that in Eq. (8), we are primarily interested in the case 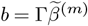.

The matrices Γ and Λ also partition the score function as 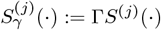 and 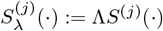. Then, the estimated (restricted) nuisance parameters 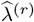 solve the restricted score equation

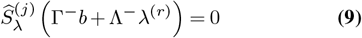

where *M* ^−^ denotes the Moore-Penrose generalized inverse of the matrix *M*. The estimated variance-covariance matrix of 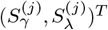 is

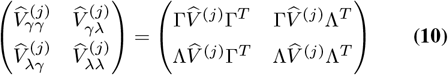

so that a score test of *H*_0_ : *β*_*j*_ = *β*^(*m*)^ can be constructed as

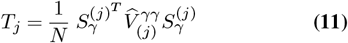

where

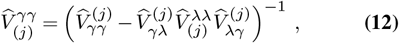

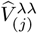 is the inverse of 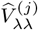 and where it is understood all quantities are evaluated at 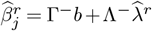. We can expect *T*_*j*_ to asymptotically have a 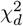 distribution with *d* degrees of freedom. If Γ is a square, full-rank matrix, then the null hypothesis is simple: Λ = 0, there are no nuisance parameters, 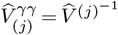 in Equation Eq. (11), and the resulting test has *d* = *K* degrees of freedom.

## Simulations

We assessed the performance of the proposed method through simulation studies. First, we evaluated the validity of the proposed variance-covariance estimator Eq. (6) and Eq. (7) by generating data according to equation Eq. (1). Because these results relate to the non-compositional AFT, descriptions of these simulations are presented in the Supplementary Section S2.

To evaluate the compositional AFT (CAFT), we generated entire microbial communities using the microbiome data simulator (MIDASim), which has been shown to produce realistic microbial communities (23), to generate datasets that capture key features of real microbiome data, including taxon-taxon correlations, sparsity, and dispersion. We used data from the Inflammatory Bowel Disease Multi-omics Database (IBDMDB) project (24) as the template for the simulated data. This dataset, sourced from the `HMP2Data` R package, comprises 178 samples and 982 taxa, with 89.69% of the cells having zero counts. For quality control, we excluded samples with library sizes below 3,000 and taxa that were absent in all samples. After this filtering, the feature table comprises 146 samples and 614 taxa, with 85.09% of the cells still containing zero counts. To compare CAFT with existing methods, we analyzed the simulated datasets using LinDA (25), ANCOM-BC2 (two versions) (26), LOCOM (3), and LDM-clr (25). For each method, we evaluated the type I error, false discovery rate (FDR), and sensitivity.

We simulated two covariates, *x*_1_ and *x*_2_, with *x*_1_ as the primary variable of interest and *x*_2_ as a variable we wish to adjust for. Both *x*_1_ and *x*_2_ were simulated as binary random variables. We implemented a balanced design, allocating one quarter of the data to each possible combination of *x*_1_ and *x*_2_ values within the set of {0,1}. We used the parametric version of MIDASim to simulate microbiome communities. Using MIDASim, we first generated mean relative abundance vector 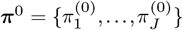 estimated from the template IBD data. We modified the relative abundance for taxon *j* as

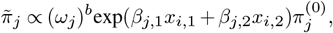

where *β*_*j*,1_ and *β*_*j*,2_ are the effect sizes associated with *x*_1_ and *x*_2_, *ω*_*j*_ is a taxon-specific bias factor, and *b* is a parameter that determines the overall effect of bias. We generated *ω*_*j*_ from a continuous uniform distribution ranging from 2 to 10, and varied the bias parameter *b* within the set {0, 1, 2}. Note that *b* = 0 corresponds to no bias in the simulation, while *b* ≠ 0 effectively alters the values of 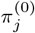 to 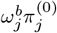 to introduce bias; as *ω*_*j*_ changes for each simulation replicate, bias has the effect of changing the baseline taxon frequencies of every taxon in each simulation replicate.

We designated *m* taxa to be ‘causal’ with *m* = 10, 20, 30. Of the causal taxa, five were associated with both *x*_1_ and *x*_2_ (*β*_*j*,1_ ≠ 0, *β*_*j*,2_ ≠ 0), five were solely associated with *x*_2_ (*β*_*j*,1_ = 0, *β*_*j*,2_ ≠ 0), and the remaining *m* − 10 taxa were solely associated with *x*_1_ (*β*_*j*,1_ ≠ 0, *β*_*j*,2_ = 0). These *m* causal taxa were randomly selected from the 50 taxa with the highest abundance level in the template data. The remaining taxa had *β*_*j*,1_ = 0 and *β*_*j*,2_ = 0. For the causal taxa, for each simulated dataset, *β*_*j*,2_ = 1 for all taxa associated with *x*_2_, and a single value for *β*_*j*,1_ was selected from the set {0.5, 1, 1.5, 2} and applied to all taxa associated with *x*_1_. The library size for each sample was determined by sampling with replacement from the library sizes in the original template data. For our simulations, we considered sample sizes of *n* = 100, 200, and 500.

We chose identifying taxa associated with *x*_1_, while adjusting for *x*_2_, as the goal of the analysis. For each simulated dataset, we attempted this goal using CAFT as well as several existing approaches: LOCOM, LinDA, ANCOM-BC2, and its robust variant, ANCOM-BC2-Robust, as well as LDM-clr. We considered two levels of filtering to remove taxa with very sparse data. In the first, we removed taxa that were present in fewer than 6% of the samples; in the second, we removed taxa present in fewer than 20% of the samples. For each analysis, we used 200 simulation replicates. The type I error was assessed by computing the average proportion of taxa incorrectly detected as associated with *x*_1_. We used the Benjamini-Hochberg procedure (27) to adjust individual p-values for FDR control.

Figure S2 presents the simulation outcomes using the 6% sample presence cutoff for taxon filtering, with *n* = 100, and in conditions with no bias. ANCOM-BC2 exhibited inflated type I error rates and poor FDR control across all scenarios, including when no taxa were associated with *x*_1_ (*β*_1_ = 0). ANCOM-BC2-Robust reported well-controlled FDR; note that ANCOM-BC2-Robust and ANCOM-BC2 use the same p-values, but differ in their FDR adjustment. LDM-clr and LinDA showed increasing type I error with larger values of *β*_1_ or *m*, leading to corresponding increases in FDR. In contrast, LOCOM and the proposed CAFT methods exhibit well-controlled type I errors. LOCOM showed a modest increase in FDR with increasing *β*_1_ at a target FDR control of 0.05. Meanwhile, the proposed CAFT method consistently controlled type I error effectively. Due to the inadequate FDR control of other methods, we restrict sensitivity evaluations to ANCOM-BC2-Robust and CAFT, noting that ANCOM-BC2-Robust has substantially lower sensitivity than CAFT. Similar results were obtained when bias was introduced (*b* = 1 or 2; see Figures S1 and S2). Furthermore, simulations with *n* = 200 or *n* = 500 showed similar results and are omitted for brevity.

## Analysis of Microbiome Data

### Gut microbiome in inflammatory bowel diseases

We analyzed gut microbiome data using a subset of 16S rRNA sequencing data from the IBDMDB project (28), the template data in our simulation studies. The original IBDMDB dataset included 178 microbiome samples from 81 patients collected across multiple visits. For this analysis, we analyzed a total of 47 samples, including 22 individuals without IBD (non-IBD), and 25 had Crohn’s disease (CD). We restricted our analysis to specimens collected at the first visit and from the ileum biopsy. Samples with library size ≤ 3000 were excluded. We further filtered out rare taxa that are present in fewer than 10% of samples, resulting in 211 taxa in the analysis. After filtering, the proportion of zeros was 65.6%. The primary variable of interest was disease status (non-IBD VS. CD) with gender as a covariate. We analyzed the data using CAFT, LOCOM, LinDA, ANCOM-BC2, ANCOM-BC2-Robust, and LDM-clr, using a nominal FDR threshold of 20%. Both covariates were centered prior to analysis.

Figure 2 displays the UpSet plot (29) summarizing the overlap of differentially abundant taxa identified by all methods. The horizontal bars represent the total number of discoveries by each method. The vertical bars indicate the number of taxa identified uniquely or jointly by different method combinations (indicated by black dots at the bottom). ANCOM-BC2 detected the largest number of differentially abundant taxa (118), of which 101 were not identified by any of the other methods. However, given the inflated type I error and elevated FDR for ANCOM-BC2 in the simulation studies, many of these taxa may be false positives. LinDA and LDM-clr reported a similar number of discoveries (23 and 19, respectively), with 15 taxa overlapping. CAFT detected 6 differentially abundant taxa. Among them, 2 were detected by all methods evaluated, 2 were detected by all methods except LOCOM, and 1 was detected by all methods except LOCOM and ANCOM-BC2-Robust, and one was not detected by any of the other methods except LinDA.

**Fig. 1.**
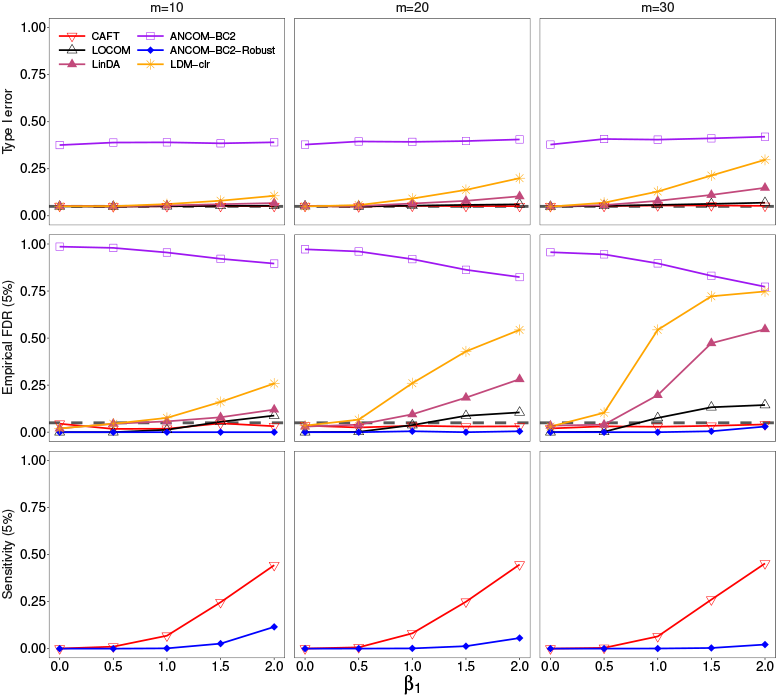
Results from the MIDASim simulation: *x*_1_ and *x*_2_ are both binary, no bias (*b* = 0), taxa filtered at 6%, n = 100. The gray dashed line indicates the nominal level Type I error of 0.05 in the first row and the FDR = 0.05 in the second row. Numbers in parentheses of row names represent the FDR cutoffs applied during the evaluation.

**Fig. 2.**
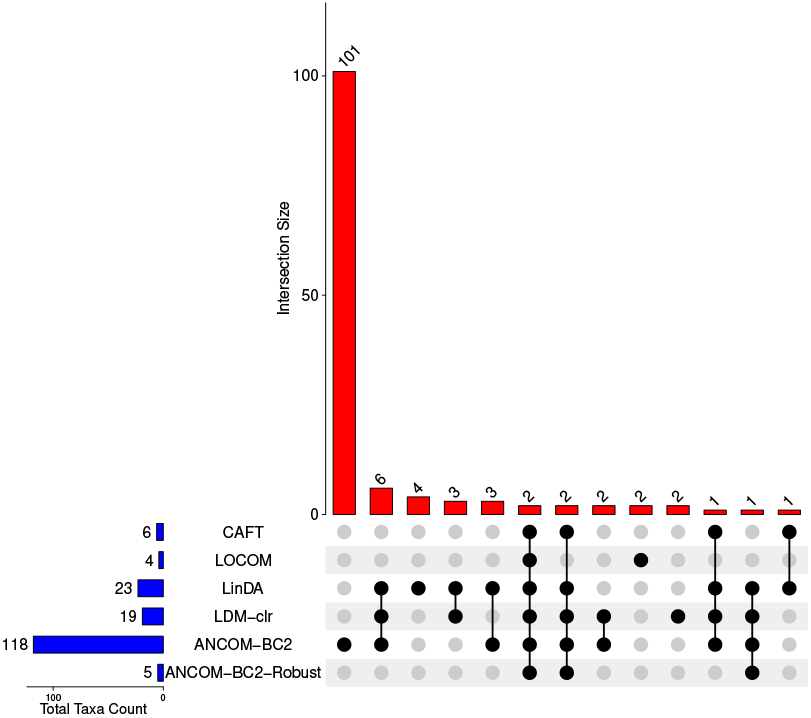
The UpSet plot illustrating the number of taxa identified by individual methods and their intersections in the IBD microbiome study, based on 10% taxon presence filter and 20% FDR. The vertical bar represents the number of taxa uniquely or jointly detected by the indicated combination of methods. Horizontal bars show the total number of taxa detected by each method.

Figure 3 presents boxplots showing the relative abundance distributions of the six taxa detected by CAFT, a representative null taxon from CAFT (the taxon corresponding to the median value of the 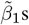), and two additional taxa that were detected by LOCOM but not by CAFT. The six CAFT-detected taxa all exhibited higher relative abundance in non-IBD participants compared to participants with CD, and this pattern was consistent in both males and females. This is in contrast to the null taxon for which the relative abundances are similar between the non-IBD and CD participants. The two taxa that were detected by LOCOM but not by CAFT were both borderline-significant using CAFT, with p-values 0.0176 (*Bacteroidaceae Bacteroides*) and 0.0134 (*Family XI Peptoniphilus*). However, neither passed the multiple comparison correction (both had BH-adjusted p-values of 0.277). The six taxa detected by CAFT are likely to have biological relevance in CD, as many belong to families such as *Lachnospiraceae* and *Ruminococcaceae*, which are known producers of short-chain fatty acids and are frequently depleted in inflammatory conditions (30, 31). Several of these taxa, including *Coprococcus, Lachnospira*, and *Eubacterium eligens group*, have been previously linked to gut barrier function and immune regulation in the context of IBD (30, 32, 33). *Lachnospiraceae UCG001* has also shown protective associations in Mendelian randomization studies (34).

**Fig. 3.**
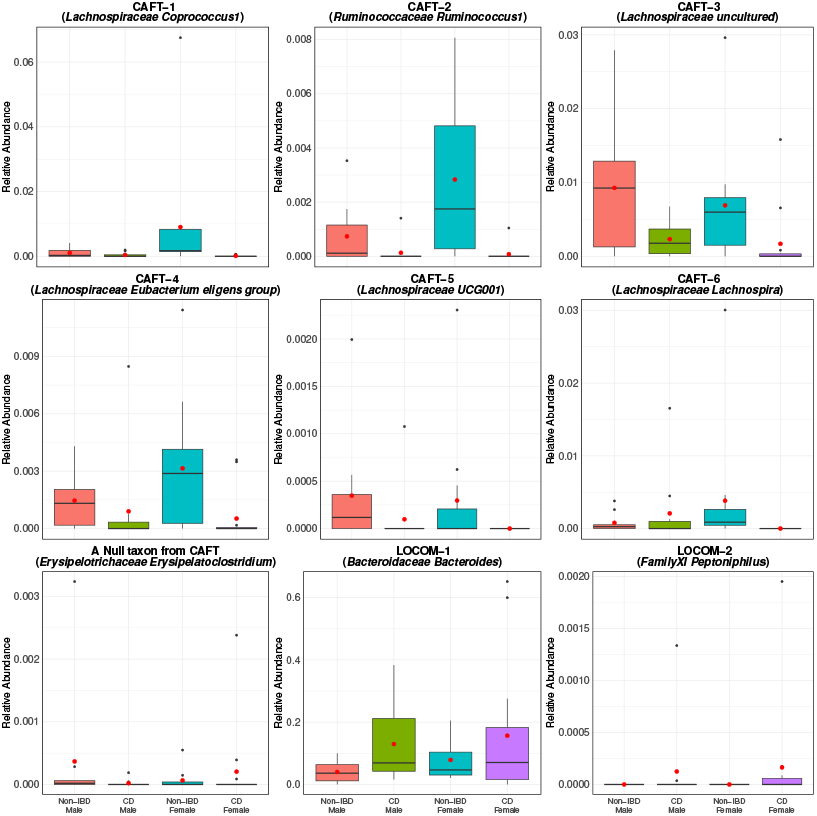
Distribution of relative abundances for several selected taxa in the IBD dataset. Red dots represent means. CAFT–1 and CAFT-2 taxa were detected by all methods. CAFT-3 and CAFT-4 taxa were detected by all other methods except LOCOM. CAFT-5 taxon were detected CAFT, LinDA, LDM-clr, and ANCOMMBC2. CAFT-6 was only detected by CAFT and LinDA. The null taxon corresponds to the taxon (*Erysipelotrichaceae Erysipelatoclostridium*) with the median of estimated 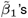 from CAFT. The subsequent two plots show two taxa that were uniquely detected by LOCOM.

### Upper respiratory tract (URT) microbiome data set

Our second dataset originates from a study investigating the effect of cigarette smoking on the oropharyngeal and nasopharyngeal microbiome (35), which was also analyzed in the LOCOM paper (3). Following (3), we focused our analysis on the microbiome data from left oropharyngeal samples, also excluded three samples from individuals who had used antibiotics in the three months prior to sample collection, and followed the same filtering steps except that we removed taxa that were present in fewer than than 10% (rather than 20% in LOCOM paper) of the samples. After these steps, the final dataset consisted of 57 samples (26 smokers and 31 non-smokers; 20 females and 37 males) and 193 taxa. After filtering, 77.5% of the OTU table consisted of zeros. We applied the CAFT approach along with competing methods to investigate the association between each taxon and smoking status, adjusting for sex. Deviating slightly from the analysis in (3), we centered the covariates prior to analysis to ensure consistency across all methods.

Figure 4 summarizes the analysis results for all methods in an UpSet plot (29), using a 10% FDR threshold. ANCOMBC2 consistently reported the most taxa (54), followed by LDM-clr (12), LinDA (9), CAFT (7), and LOCOM (2), while ANCOM-BC2-Robust remained highly conservative and detected none. Similar to our analysis of the IBD dataset, the large number of discoveries by ANCOM-BC2 may reflect an inflated false positive rate. Notably, every taxon detected by CAFT and LOCOM was also identified by at least two other methods, suggesting the robustness of these findings. Compared to LOCOM, CAFT identified five additional taxa.

**Fig. 4.**
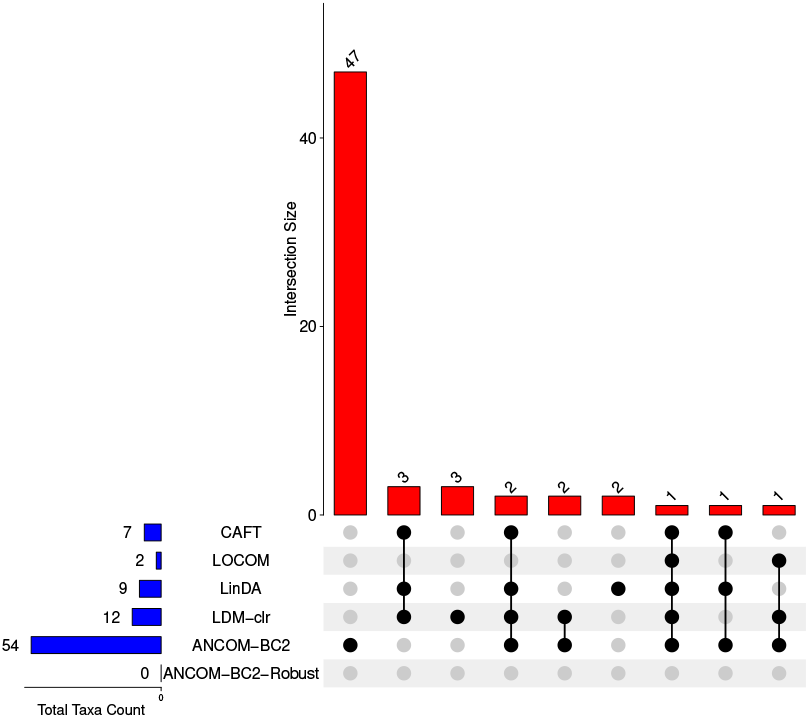
The UpSet plot illustrating the number of taxa identified by individual methods and their intersections using 10% presence filter and 10% FDR in the URT microbiome study.

In Figure 5, we present the boxplot of the relative abundances of the seven CAFT-detected taxa, together with a representative null taxon (corresponding to the taxon having the median of the estimated 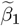 values), and one taxon that was detected by LOCOM but not CAFT. Among them, all but *Veillonellaceae Veillonella* (labeled CAFT-3 in the figure) show higher relative abundance in smokers than non-smokers. Note this is not the case in the null taxon (*Veillonellaceae Dialister*). The taxon detected by LOCOM but not by CAFT, *Prevotellaceae Prevotella* (labeled LOCOM-1 in the figure), showed borderline significance under CAFT with an unadjusted p-value of 0.0127, but did not remain significant after multiple comparison correction (BH-adjusted p-value 0.163).

**Fig. 5.**
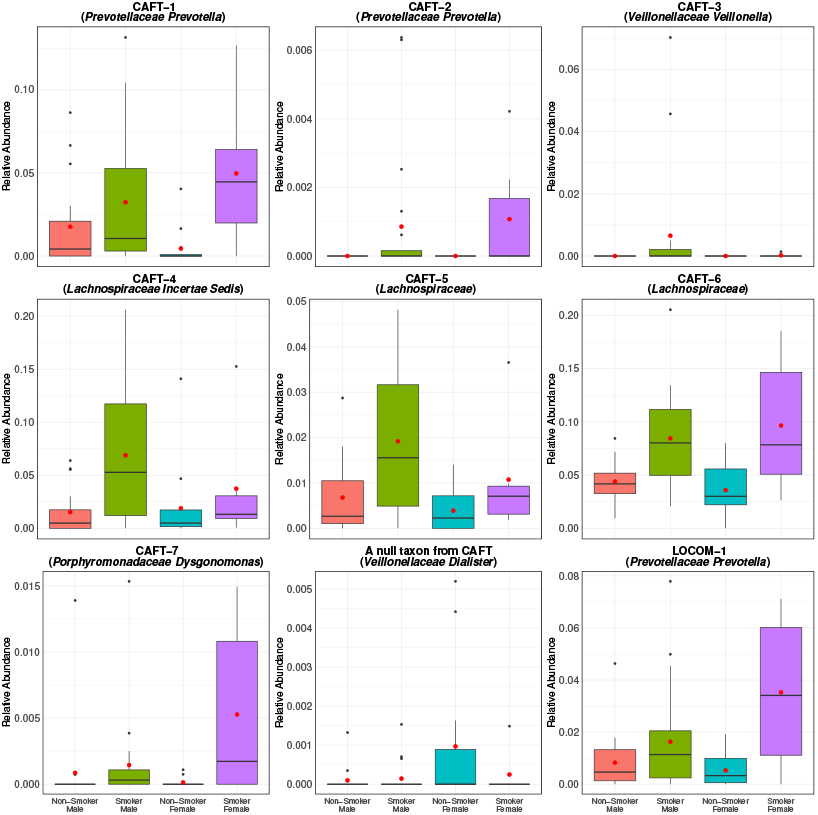
Distribution of relative abundances for several selected taxa in URI data. Red dots represent the means. CAFT-1, CAFT-2, and CAFT-3 were detected by CAFT, LinDA, and LDM-clr. CAFT-4 and CAFT-5 taxa were detected by CAFT, LinDA, LDM-clr, and ANCOM-BC2. CAFT-6 was detected by CAFT, LOCOM, LinDA, LDM-clr, and ANCOM-BC2. CAFT-7 was detected by CAFT, LinDA, LDM-clr, and ANCOMBC2. The null taxon corresponds to the taxon (*Veillonellaceae Dialister*) with the median of estimated 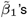 from CAFT. The last plot shows a taxon that was uniquely detected by LOCOM, LDM-clr, and ANCOM-BC2.

The CAFT-detected taxa show smoking-associated patterns consistent with prior reports. *Veillonella* is more enriched in smokers compared to nonsmokers (35), and a shotgun metagenomics study similarly reported higher *Prevotella* in smokers (36), in line with our observations. Smokers also showed enrichment of anaerobic lineages in the upper respiratory tract; members of the family *Lachnospiraceae* were increased in smokers’ airways (35). *Porphyromonadaceae Dysgonomonas* shows higher relative abundance in smoke groups and has been reported in smoker-linked oral airway conditions (37). As a representative null taxon, *Veillonellaceae Dialister*, was not identified as differentially abundant by either method and displayed flat patterns across groups, and served as a visual reference for non-associated taxa. Finally, *Prevotellaceae Prevotella*, identified by LOCOM, LDM-clr, and ANCOM-BC2, exhibits a clear differential pattern in which the smoker-female subgroup shows the highest relative abundance among the four groups.

## Discussion

In this study, we developed a compositional approach to identify microbial taxa associated with specific variables, with adjustments for covariates. Microbiome data are characterized by high sparsity, often with more than 50% of cells exhibiting zero read counts. Traditional compositional analysis methods for microbiome data, such as LinDA (6), LDM-clr (7), ANCOM-BC2 (9), and the robust version of ANCOM-BC2, typically rely on pseudocounts. Only LOCOM avoids pseudocounts by fitting a series of logistic regressions. In contrast, our method interprets zero counts as censored observations within a survival analysis framework, utilizing the AFT model, thereby circumventing the need for pseudocounts. We note here that Microbiome data are compositional because the library size *N*_*i*_ is not informative of the microbial load in a specimen, but rather reflects technical aspects such as the efficiency of the PCR step. Thus, it is worth noting that treating 1*/N*_*i*_ as a limit of detection is consistent with the information conveyed by the library size.

The performance of survival analysis models frequently degrades as the proportion of censoring increases. In microbiome data, the proportion of censored observations can be quite large. Our choice of the (16) and (17) estimator in Eq. (3) and Eq. (4) represents the result of an extensive search for an estimation method for the AFT that performed well at the levels of censoring found in microbiome data. Further, the improved performance of our empirical variance estimator Eq. (6) and Eq. (7) over the standard Cox-model-based variance estimator of (19) also makes the asymptotic inference in CAFT reliable, as demonstrated by our simulation studies.

Converting the log-linear models for relative abundance at each of *J* taxa, which yields *J* regression coefficients 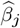, into a compositional model requires reducing the number of 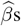 by one. This can be accomplished in a variety of ways, e.g., by applying the constraint that 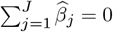 as in (4) and (5). Here, we adopt the approach first proposed in (3), in which we assume the large majority of taxa have the null effect, so that the median 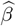, denoted 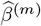, should correspond to a null taxon. Then, we base inference on 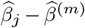. We note that LinDA (6) uses the modal 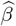, while LDM-clr (7) offers both options. Although either of these assumptions is typically realistic, they may not always hold, particularly in analyses at higher taxonomic levels, such as class or phylum, where a significant proportion of taxa may be associated with the variable of interest. Additionally, CAFT treats 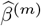, which is estimated from data at all taxa, as a fixed parameter in hypothesis tests at each taxon. While this simplification is justified when the number of taxa is large, it becomes problematic with datasets having a small number of taxa. We also note that estimating the modal 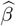 becomes more difficult as the number of components in *β* increases, while several multivariate estimators of median are available (CAFT uses the affine equivariant multivariate median of (22)).

Because CAFT uses asymptotic variance estimators, it is computationally efficient. This advantage is particularly notable for datasets with larger sample sizes and a higher number of taxa. As shown in Table 1, CAFT consistently required only a few seconds to complete the analysis across all scenarios. This efficiency is largely due to its improved variance estimator, which eliminates the need for computationally intensive permutation or resampling procedures. In contrast, LOCOM—which relies on permutation-based inference, exhibited substantially longer runtimes, especially under the 10% presence filter, taking up to one minute for the IBD data. While LinDA achieved the fastest runtimes overall, its lack of robust error control limits its reliability, as previously discussed. LDM-clr and ANCOM-BC2 displayed moderate computation times, falling between CAFT and LOCOM.

**Table 1.** Computation Time (in seconds) for the comparing methods on IBD and URT data. All methods were run in a single thread on a Mac Studio with an M2 Max CPU and 64 GB of memory.

| Methods | IBD | URT |
| --- | --- | --- |
| CAFT | 4.081 | 4.585 |
| LOCOM | 60.291 | 43.812 |
| LinDA | 0.037 | 0.036 |
| LDM-clr | 15.244 | 4.927 |
| ANCOM-BC2 | 14.741 | 15.888 |

## Author contributions statement

GS and ML wrote the R code for the CAFT package. ML and NZ contributed to the development and testing of the methods, analyzed the data, and wrote the manuscript. GS developed the method and wrote the manuscript. All authors read the manuscript and approved the final manuscript.

## Data Availability Statement

The IBD and URI datasets are distributed with the MIDASim and HMP2Data R packages, respectively, and can be obtained from https://doi.org/doi:10.18129/B9.bioc.HMP2Data and https://github.com/yijuanhu/LOCOM.

## S1: The Cox variance estimator

The Cox variance estimator based on derives from a Martingale representation of the Cox model. Dropping the superscript for taxon (*j*), the Martingale representation of *S* is

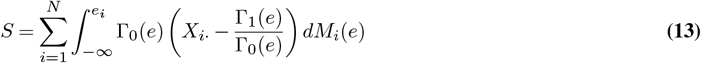

where

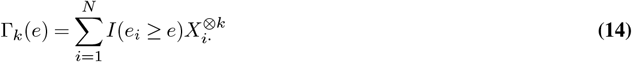

and *dM*_*i*_(*e*) = *dN*_*i*_(*e*) − *Y* (*e*)*d*Λ(*e*) and *N*_*i*_(*e*) = *I*(*e*_*i*_ ≤ *e*), where Λ is the cumulative hazard and *Y* is the at-risk indicator. Then, *S* is a weighted version of the Cox score with weight Γ_0_(*e*). We then define 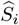 as

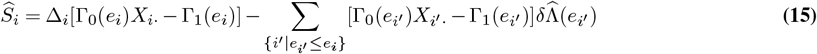

where 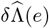 is the jump in the estimated cumulative hazard Λ at residual *e*. Then, the Cox variance proposed by Zeng and Lin (19) is

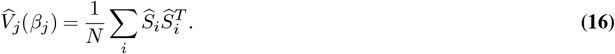

## S2: Additional Simulations

To assess the performance of our variance estimators for the AFT model, we simulated survival time using the following equation,

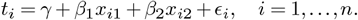

Here, *x*_*i*1_ was the variable of interest and *x*_*i*2_ served as the adjustment variable. We generated *x*_*i*1_ from a standard normal distribution and induced correlations between covariates via 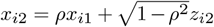 with *z*_*i*2_ ∼ *N* (0, 1). The correlation parameter *ρ* ∈{−0.5, 0, 0.3, 0.5} controls the direction and strength of the confounding effect. Error terms *E*_*i*_ were generated independently from four distributions: *N* (0, 1), Exponential(1), Cauchy(0,1) and Weibull(*k* = 1.5, *λ* = 1), each centered to have mean zero. The censoring time *c*_*i*_ was drawn as *c*_*i*_ ∼ *N* (*µ*_*c*_, 1). The observed survival time was *τ*_*i*_ = min(*t*_*i*_, *c*_*i*_) with censoring indicator Δ_*i*_ = *I*(*t*_*i*_ *< c*_*i*_). We considered two censoring scenarios by choosing *µ*_*c*_ = − 0.9 to yield moderate censoring (around 60-70%, Table S1) and choosing *µ*_*c*_ = −3.3 to yield high censoring (around 80-98%, Table S2). The detailed average censoring proportions for different error distributions appear in the tables. For type-I error evaluation, we fixed *β*_1_ = 0, *β*_2_ = 1, and *γ* = 0. Each design was replicated 10,000 times with *n* ∈ {100, 200, 500}. In each replicate, we tested *H*_0_ : *β*_1_ = 0 using the proposed CAFT variance estimator for the AFT model and, for comparison, a COX variance estimator, and recorded the empirical rejection proportion at *α* = 0.05.

**Table S1.** Empirical type-I error (*α* = 0.05) for testing *H*_0_ : *β*_1_ = 0 using the proposed CAFT variance–covariance estimator for the AFT model and a Cox-based variance estimator (COX). Results are shown by error distribution (Normal, Exponential, Cauchy, Weibull), sample size (*n* = 100, 200, 500), confounding correlation *ρ* ∈ *{*−0.5, 0, 0.3, 0.5*}*, and the moderate average censoring proportion (around 60 - 70 %). Each entry is the rejection proportion over 10,000 simulations

| Distribution | Censoring Proportion | $n$ | Method | $\rho$ | | | |
| --- | --- | --- | --- | --- | --- | --- | --- |
|  |  |  |  | -0.5 | 0 | 0.3 | 0.5 |
| Normal | ~ 70% | 100 | CAFT | 0.0482 | 0.0458 | 0.0481 | 0.0498 |
|  |  |  | COX | 0.0566 | 0.0543 | 0.0529 | 0.0569 |
|  |  | 200 | CAFT | 0.0490 | 0.0501 | 0.0486 | 0.0524 |
|  |  |  | COX | 0.0530 | 0.0518 | 0.0536 | 0.0573 |
|  |  | 500 | CAFT | 0.0508 | 0.0528 | 0.0498 | 0.0517 |
|  |  |  | COX | 0.0539 | 0.0525 | 0.0506 | 0.0524 |
| Exponential | ~ 70% | 100 | CAFT | 0.0496 | 0.0441 | 0.0509 | 0.0493 |
|  |  |  | COX | 0.0584 | 0.0566 | 0.0571 | 0.0586 |
|  |  | 200 | CAFT | 0.0476 | 0.0487 | 0.0487 | 0.0506 |
|  |  |  | COX | 0.0543 | 0.0538 | 0.0521 | 0.0533 |
|  |  | 500 | CAFT | 0.0483 | 0.0485 | 0.0481 | 0.0456 |
|  |  |  | COX | 0.0506 | 0.0500 | 0.0486 | 0.0471 |
| Cauchy | ~ 60% | 100 | CAFT | 0.0535 | 0.0496 | 0.0478 | 0.0504 |
|  |  |  | COX | 0.0650 | 0.0646 | 0.0647 | 0.0673 |
|  |  | 200 | CAFT | 0.0500 | 0.0491 | 0.0508 | 0.0509 |
|  |  |  | COX | 0.0586 | 0.0577 | 0.0589 | 0.0589 |
|  |  | 500 | CAFT | 0.0503 | 0.0529 | 0.0533 | 0.0515 |
|  |  |  | COX | 0.0526 | 0.0549 | 0.0557 | 0.0531 |
| Weibull | ~ 72% | 100 | CAFT | 0.0451 | 0.0492 | 0.0459 | 0.0477 |
|  |  |  | COX | 0.0568 | 0.0598 | 0.0582 | 0.0561 |
|  |  | 200 | CAFT | 0.0510 | 0.0524 | 0.0558 | 0.0524 |
|  |  |  | COX | 0.0567 | 0.0584 | 0.0603 | 0.0571 |
|  |  | 500 | CAFT | 0.0470 | 0.0511 | 0.0517 | 0.0484 |
|  |  |  | COX | 0.0477 | 0.0540 | 0.0541 | 0.0513 |

Across all designs, the proposed CAFT variance estimator maintained nominal type-I error, while the COX estimator was increasingly liberal as censoring intensified. Under moderate censoring (around 60 - 70 % in Table S1), CAFT stayed close to 0.05 for every error distribution and *n* = 100, 200, 500, including the heavy-tailed Cauchy case. COX was generally near nominal but tended to over-reject, most noticeably with Cauchy errors. Under high censoring (around 80 - 98 % in Table S2), CAFT again hovered around 0.05 across scenarios, roughly from 0.047 to 0.052, whereas COX showed marked inflation for Exponential and Weibull errors, about 0.12 to 0.16 at *n* = 100 and 0.08 to 0.13 at *n* = 200. For *n* = 500, the type I error inflation reduced (about 0.05 to 0.06) but is still above CAFT. For Normal errors, COX remained near nominal under high censoring, and for Cauchy, it was slightly inflated when the sample size is small (*n* = 100 and 200). Variations across the correlation parameter *ρ* were minor relative to the effects of censoring level, error distribution, and sample size.

**Table S2.** Empirical type-I error (α = 0.05) for testing *H*_0_ : β_1_ = 0 using the proposed CAFT variance–covariance estimator for the AFT model and a Cox-based variance estimator (COX). Results are shown by error distribution (Normal, Exponential, Cauchy, Weibull), sample size (n = 100, 200, 500), confounding correlation *ρ* ∈ {−0.5, 0, 0.3, 0.5}, and the high average censoring proportion (around 80 - 98 %). Each entry is the rejection proportion over 10,000 simulations.

| Distribution | Censoring Proportion | $n$ | Method | $\rho$ | | | |
| --- | --- | --- | --- | --- | --- | --- | --- |
|  |  |  |  | -0.5 | 0 | 0.3 | 0.5 |
| Normal | $\sim 97\%$ | 100 | CAFT | 0.0493 | 0.0501 | 0.0475 | 0.0466 |
|  |  |  | COX | 0.0552 | 0.0519 | 0.0537 | 0.0572 |
|  |  | 200 | CAFT | 0.0490 | 0.0503 | 0.0492 | 0.0501 |
|  |  |  | COX | 0.0286 | 0.0320 | 0.0288 | 0.0312 |
|  |  | 500 | CAFT | 0.0524 | 0.0502 | 0.0488 | 0.0497 |
|  |  |  | COX | 0.0495 | 0.0523 | 0.0515 | 0.0501 |
| Exponential | $\sim 98\%$ | 100 | CAFT | 0.0479 | 0.0404 | 0.0429 | 0.0467 |
|  |  |  | COX | 0.1516 | 0.1390 | 0.1401 | 0.1504 |
|  |  | 200 | CAFT | 0.0520 | 0.0506 | 0.0474 | 0.0515 |
|  |  |  | COX | 0.0883 | 0.0808 | 0.0887 | 0.0890 |
|  |  | 500 | CAFT | 0.0508 | 0.0506 | 0.0527 | 0.0482 |
|  |  |  | COX | 0.0556 | 0.0479 | 0.0593 | 0.0525 |
| Cauchy | $\sim 81\%$ | 100 | CAFT | 0.0457 | 0.0471 | 0.0474 | 0.0474 |
|  |  |  | COX | 0.0674 | 0.0661 | 0.0666 | 0.0720 |
|  |  | 200 | CAFT | 0.0483 | 0.0493 | 0.0479 | 0.0490 |
|  |  |  | COX | 0.0617 | 0.0609 | 0.0600 | 0.0589 |
|  |  | 500 | CAFT | 0.0485 | 0.0487 | 0.0483 | 0.0490 |
|  |  |  | COX | 0.0557 | 0.0541 | 0.0543 | 0.0537 |
| Weibull | $\sim 98\%$ | 100 | CAFT | 0.0401 | 0.0357 | 0.0377 | 0.0437 |
|  |  |  | COX | 0.1562 | 0.1424 | 0.1430 | 0.1532 |
|  |  | 200 | CAFT | 0.0494 | 0.0450 | 0.0413 | 0.0473 |
|  |  |  | COX | 0.1224 | 0.1211 | 0.1184 | 0.1286 |
|  |  | 500 | CAFT | 0.0511 | 0.0475 | 0.0485 | 0.0512 |
|  |  |  | COX | 0.0554 | 0.0494 | 0.0521 | 0.0535 |

## S3: Supplementary Figures

**Fig. S1.**
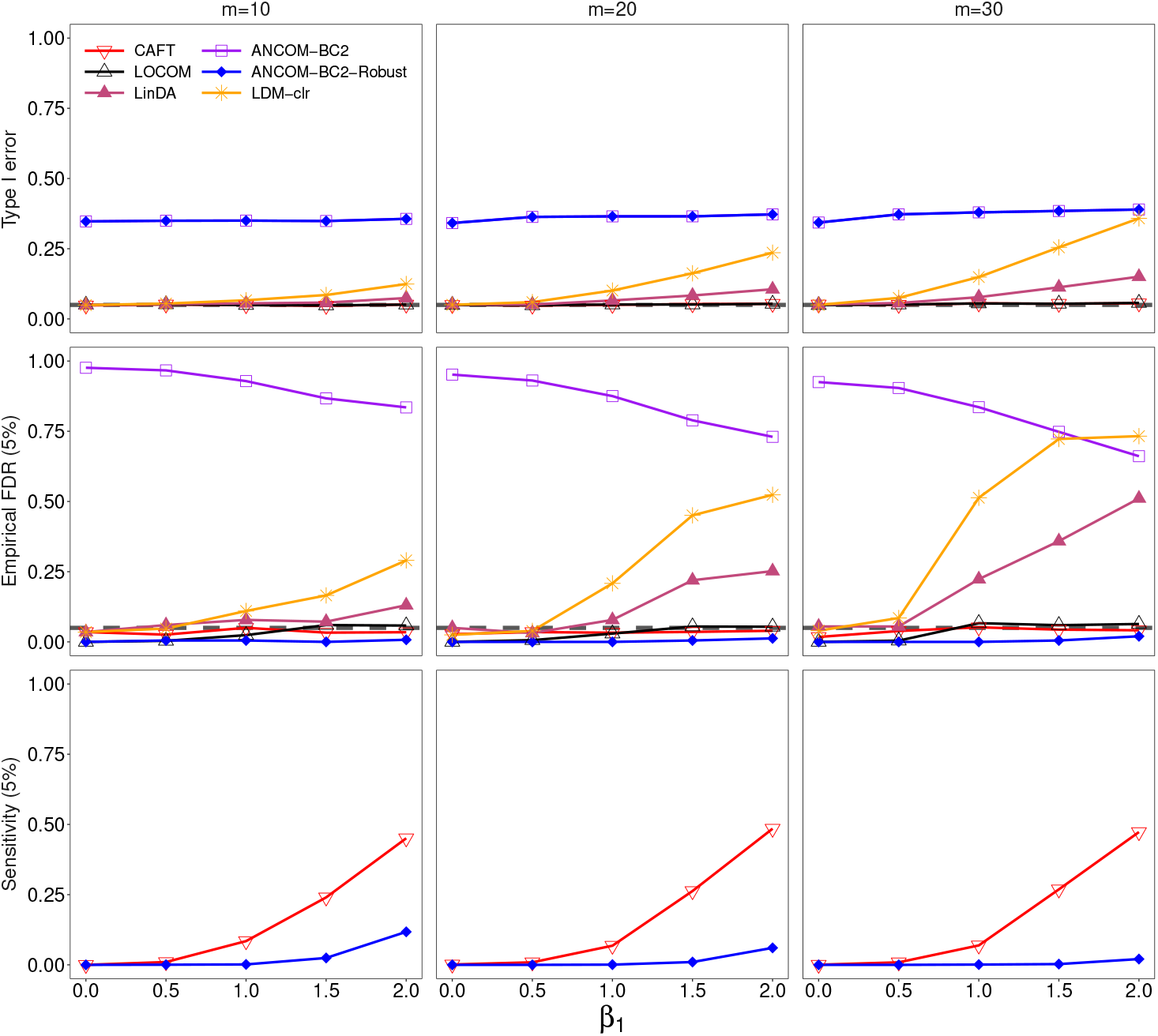
Results from the MIDASim simulation: *x*_1_ and *x*_2_ are both binary, bias (*b* = 1), taxa filtered at 6%, n = 100. The gray dashed line indicates the nominal level Type I error of 0.05 in the first row. Numbers in parentheses of row names represent the FDR cutoffs applied during the evaluation.

**Fig. S2.**
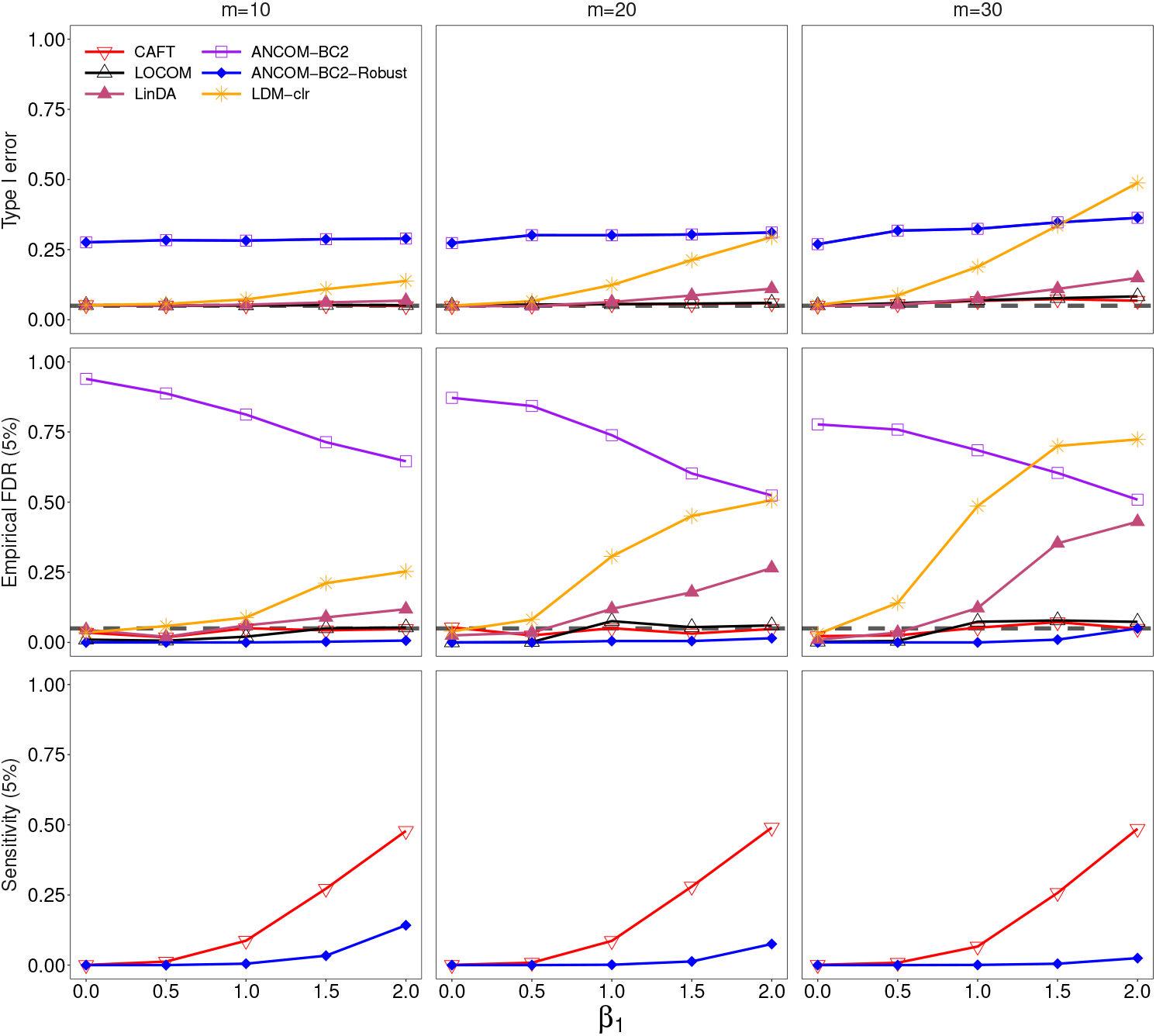
Results from the MIDASim simulation: *x*_1_ and *x*_2_ are both binary, bias (*b* = 2), taxa filtered at 6%, n = 100. The gray dashed line indicates the nominal level Type I error of 0.05 in the first row. Numbers in parentheses of row names represent the FDR cutoffs applied during the evaluation.

